# Preclinical Study: Sunitinib-suppressed MiR-452-5p Facilitates Renal Cancer Cell Invasion and Metastasis Through Modulating SMAD4/SMAD7/EMT Signals

**DOI:** 10.1101/286898

**Authors:** Saiyang Li, Jin Zhang, Yonghui Chen, Junjie Ma, Wen Kong, Dongkui Gong, Junhua Zheng, Wei Xue, Wei Zhai, Yunfei Xu

## Abstract

Although microRNAs (miRNAs) have been revealed as crucial modulators in tumor metastasis and target therapy, our understanding of their roles in metastatic renal cell carcinoma (mRCC) and Sunitinib treatment is limited. Here, We focused on 2 published microarray data to select out our anchored miRNA which was downregulated after Sunitinib treatment while upregulated in metastasis RCC tissues. Then we discovered that treating with Sunitinib, the targeted receptor tyrosine kinase inhibitor (TKI), inhibited renal cell migration and invasion via attenuating the expression of miR-452-5p. The novel identified miR-452-5p was upregulated and associated with poor prognosis in RCC. Preclinical studies using multiple RCC cells and xenografts model illustrated that miR-452-5p could promote RCC cell migration and invasion *in vitro* and *in vivo*. Mechanistically, P65 could directly bind to the miR-452-5p promoter and thus transcriptionally induce miR-452-5p expression, which led to post-transcriptionally abrogate SMAD4 expression, thus inhibition of its downstream signals including SMAD7 and EMT (Epithelial-Mesenchymal Transition) associated genes. Our study presented a road map for targeting this newly identified miR-452-5p and its SMAD4/SMAD7/EMT signals pathway, which imparted a new potential therapeutic strategy for mRCC treatment.

## Introduction

Renal cell carcinoma (RCC) is one of the most aggressive human urological malignancies, accounting for 2%–3% of adult malignancies [15; 39]. One third of patients have metastatic lesions detected at primary diagnosis, and 30% eventually develop metastatic renal cell carcinoma (mRCC) after surgery [2]. Because of its resistance to radiotherapy and chemo-therapy, surgical resection remains the unique effective treatment [1; 25]. Hence, a better understanding of the detailed mechanisms behind the pathogenesis of RCC and more effective treatment strategies are urgently required.

MicroRNAs (miRNAs), a group of small noncoding RNAs of about 20-24 nucleotides in length, negatively regulate gene expression [11; 30]. Accumulating evidence has demonstrated the involvement of miRNAs contributing to multiple metastatic steps in various human cancers [8; 38]. Notably, plenty of miRNAs, such as miR-27a, miR-141 and miR-205 were reported to exert their roles in RCC cell invasion and metastasis [3; 18; 22]. In addition, our group already attested that miR-646 could suppress RCC cell migration and invasion [21]. Previous studies demonstrated that miR-452-5p acted as a tumor inducer or suppressor in multiple human cancers, involving osteosarcoma, breast cancer, non-small cell lung cancer, hepatocellular carcinoma and prostate cancer [9; 12; 20; 47; 49]. Nevertheless, whether miR-452-5p is implicated in the metastasis of RCC is still need to be investigated.

Sunitinib, as the first-choice tyrosine kinase inhibitor (TKI) in mRCC, provides clinical benefits for patients with mRCC [4; 6; 27]. In a phase II study of Intermittent Sunitinib in mRCC, a total of 20 patients with a median decrease in tumor burden of 45% (range 13% to 86%) entered the intermittent phase. Most patients exhibited a stable sawtooth pattern of tumor burden (TB) reduction while receiving sunitinib and TB increase while not receiving sunitinib, and metastatic progression-free survival and overall survival were 22.4 and 34.8 months, respectively [32]. Based on this finding above, Sunitinib is recognized one of the standard therapies in mRCC. However, most of patients with mRCC will develop resistance to the drug after receiving the treatment eventually [33]. In order to predict treatment response, we urgently need promising molecular biomarkers for rational indication of TKIs in patients with mRCC. Notably, our team previously reported that Sunitinib repressed RCC progression via inducing LncRNA-SARCC, which suggested that other potential non-coding RNAs were involved in RCC-Sunitinib treatment [46]. Similarly, another preclinical study reported that the efficacy of Sunitinib could be increased by miR-32-5p [42]. Then we focused on the relationship between Sunitinib and miRNAs, which the potential ability of miRNAs to predict response to TKIs in mRCC has not drawn enough attention.

P65, as a subunit of NF-κB, targets a large number of genes and plays an important role in the regulation of apoptosis, tumorigenesis, inflammation, and various autoimmune diseases [40]. Increased P65 activation was reported to be implicated in the development of renal cell carcinoma metastasis, promoting metastasis and progression of many cancers through EMT [37]. Recently, EMT was demonstrated to be a major mechanism responsible for invasion and metastasis of cancers [31]. However, the molecular mechanism downstream of P65 has not been fully elucidated in RCC.

In our work, we demonstrated that miR-452-5p acted as a potential therapeutic target for Sunitinib and novel tumor contributor, which promoted renal cancer progression both *in vitro* and *in vivo*. Furthermore, the transactivation of miR-452-5p in RCC was induced by P65. In addition, we explored that miR-452-5p directly bind to SMAD4 and suppress SMAD4 expression, thereby regulating SMAD4/SMAD7/EMT signaling pathway and finally enhancing RCC cell invasion and metastasis. Here, we identified miR-452-5p as a novel tumor inducer as well as a potential diagnostic or prognostic marker in mRCC.

## Materials and Methods

### Tissue samples

Tumor samples and paired normal tissues from RCC patients were obtained from the Department of Urology, Shanghai Tenth People’s Hospital, Tongji University (Shanghai, China). The fresh tissues were kept in liquid nitrogen to protect the RNA from degradation. The current study was approved by the ethics committee of Shanghai Tenth People’ s Hospital.

### Cell culture and transfection

The human RCC cell lines, OSRC-2, SW839, A498, SN12-PM6 and were originally purchased from Cell Bank of the Chinese Academy of Sciences (Shanghai, China). SW839, A498 and SN12-PM6 were cultured in Dulbecco’s Modified Eagle’s Medium (DMEM, Gibco, Grand Island, New York, USA) plus 10% Fetal Bovine Serum (FBS, Hyclone, Logan, Utah, USA) with 1% penicillin/streptomycin (P/S, Gibco, Grand Island, New York, USA). OSRC-2 cells were cultured in RMPI 1640 (Gibco, Grand Island, New York, USA) plus 10% fetal bovine serum (FBS, Hyclone, Logan, Utah, USA) with 1% penicillin/streptomycin (P/S, Gibco, Grand Island, New York, USA). HK-2 cells were cultured in keratinocyte medium (KM, ScienCell, San Diego, California, USA) plus 1% keratinocyte growth supplement (KGS, Scien-Cell, San Diego, California, USA) with 1% penicillin/streptomycin (P/S, ScienCell, San Diego, California, USA). All cells described above were cultured at 37 °C in the humidified 5% CO2 environment.

### Quantitative real-time PCR (qRT-PCR)

Total RNA was extracted from tissues or cells using Trizol reagent (Invitrogen). cDNAs were synthesized with PrimeScript RT reagent Kit (Takara, Kusatsu, Japan). qRT-PCR was performed with KAPA SYBR FAST qPCR Kit (Kapa Biosystems, Woburn, Massachusetts, USA) using a 7900HT Fast Real-Time PCR System (Applied Biosystems, Carlsbad, California, USA). The primers were listed in Supplementary Table S1. The expression levels of miRNA were normalized to endogenous small nuclear RNA U6, and the expression levels of mRNA were normalized to endogenous control GAPDH. The 2^-ΔΔCt^ method was used to analyze the expression levels normalized to the endogenous control.

### Wound-healing assay and invasion assay

To analyze the migration, indicated cells were plated in six-well plates. Streaks across the plate were created in the monolayer with a pipette tip. Progression of migration was observed and photographed at 0h and 24 h after wounding. The data shown were representative micrographs of wound-healing assay of the indicated cells. The invasive capability of RCC cells was determined by the transwell assay. The membrane was coated with the matrigel (200 ng/mL; BD Biosciences). Then RCC cells were harvested and seeded with serum-free DMEM into the upper chambers at 5 × 10^4^ cells/well, and the bottom chambers contained DMEM with 10% FBS, and then transwells incubated for 24 h at 37 °C. Following incubation, the invasive cells invading into the lower surface of the membrane were fixed by 4% paraformaldehyde and stained with 1% toluidine blue. Experiments were repeated at least three times with similar results.

### Cell transfection and vector construction

According to manufacturer’s protocol, miR-452-5p/miR-NC (GenePharma, Shanghai, China), anti-miR-452-5p/anti-miR-NC (GenePharma, Shanghai, China) and LNA-miR-452-5p/LNA-miR-NC (IBS Solutions Co. Ltd, China) were transfected at a final concentration of 50 nM in OSRC-2 and SW839 cells using Lipofectamine 2000 (Invitrogen, Carlsbad, California, USA). In the rescue experiment, OSRC-2 and SW839 cells were co-transfected with 50 nM oe-SMAD4 (Lingke, Shanghai, China) and 50 nM miRNA mimics (miR-452-5p or miR-NC) with Lipofectamine 2000. The cells were harvested at 48 hr after transfection.

Lentiviral sh-miR-452-5p/sh-miR-NC was purchased from Lingke Biotechnology (Shanghai, China). To generate the stable cell lines, OSRC-2 cell line with firefly luciferase expression was transduced with the lentiviruses all above for 2 weeks. The stably knocking down cell lines were identified using qRT-PCR.

### Mouse model of orthotopic tumor implantation

OSRC-2 cells transfected with miR-452-5p, miR-452-5p+ Sunitinib, sh-miR-452-5p, sh-miR-control, oe-SMAD4 and miR-452-5p+oe-SMAD4, were injected at 2 × 10^6^ cells/mouse from ShanghaiSipper-BK laboratory animal Company (Shanghai, China), into the left subrenal capsule of 6-week-old male athymic nude mice. Cells were also transduced with luciferase *in vivo* imaging system that was performed once a week. Studies on animals were conducted with approval from the ethics committee of Shanghai Tenth People’ s Hospital.

### Chromatin immunoprecipitation assay (ChIP)

Cells were crosslinked with 4% formaldehyde for 10 min followed by cell collection and sonication with a predetermined power to yield genomic DNA fragments of 300 bp long. Lysates were precleared sequentially with normal rabbit IgG and protein A agarose. Anti-P65 antibody (2.0 μg) was added to the cell lysates and incubated at 4 °C overnight. For the negative control, IgG was used in the reaction. Specific primer sets designed to amplify a target sequence within human miR-452-5p promoter were listed in Table S1; PCR products were analyzed by agarose gel electrophoresis.

### Luciferase reporter assay

To confirm whether AR and HOXD9 could increase miR-452-5p promoter activity, OSRC-2 cells transfected with oe-P65 or vector cultured in 48-well plates were co-transfected with 1.5 μg of firefly luciferase reporter and 0.35 ng Renilla luciferase reporter with Lipofectamine 2000. Luciferase reporter assay using the one step directed cloning kit (Novoprotein, Shanghai, China) according to the manufacturer’s manual. To identify potential downstream genes of miR-452-5p, luciferase reporter assay was performed as the above described. In briefly, OSRC-2 cells transfected with miR-452-5p or vector cultured in 48-well plates were cotransfected with 1.5 μg of firefly luciferase reporter and 0.35 ng Renilla luciferase reporter with Lipofectamine 2000 regents. Luciferase reporter assay using the one step directed cloning kit.

### Protein extracting and Western blotting analysis

The cells were lysed using RIPA buffer plus protease inhibitors and phosphatase inhibitors. For western blot analysis, 25 μg of protein extracts were loaded to 10% sodium dodecylsulfate–polyacrylamide gel electrophoresis gels and transferred to nitrocellulose membranes. The membranes were incubated with a primary antibody overnight and were incubated with a secondary antibody in 1 hr with room temperature. The expression of β-actin was used as loading control. The information of antibodies were listed as follow: SMAD4(1:1000, Sangon Biotech, China, D220124), SMAD7 (1:1000, Sangon Biotech, China, D160746), E-cadherin(1:1000, Cell Signaling Technology, USA, #3195), CK-18(1:1000, Cell Signaling Technology, USA, #4548), N-cadherin(1:1000, Cell Signaling Technology, USA, #13116), Vimentin(1:1000, Cell Signaling Technology, USA, #5741).

### Immunohistochemistry

Immunohistochemistry was performed using a primary antibody against SMAD4 (1:800, Abcam, ab129108), SMAD7 (1:100, Abcam, ab131443), The degree of positivity was initially classified according to scoring both the proportion of positive staining tumor cells and the staining intensities. Scores representing the proportion of positively stained tumor cells were graded as: 0 (<10%); 1 (11–25%); 2 (26–50%); 3 (51–75%) and 4 (>75%). The intensity of staining was determined as: 0 (no staining); 1 (weak staining=light yellow); 2 (moderate staining=yellow brown); and 3 (strong staining=brown). The staining index was calculated as the product of staining intensity × percentage of positive tumor cells, resulting in scores of 0, 1, 2, 3, 4, 6, 8, 9 and 12. Only cells with clear tumor cell morphology were scored.

### Statistical analysis

Results are expressed at least 3 independent experiments. Using the GraphPad Prism statistical program, data were analyzed using ANOVA or Student’ t test unless otherwise specified. P values <0.05 were considered significant.

## Result

### Sunitinib abrogates RCC cell invasion and metastasis via depressing miR-452-5p

Our team has previously reported that Sunitinib remarkably blunted RCC progression via inducing LncRNA-SARCC [46]. In an attempt to further explore whether Sunitinib inhibited RCC cell invasion and metastasis in a miRNA-dependent manner, we first focused on 2 microarray data, GSE32099 (differentially expressed miRNAs in peripheral blood under sunitinib treatment, Table S2) and GSE37989 (metastasis-associated miRNAs, Table S3) through searching GEO datasets (Figure 1A). Next we selected out top 10 common miRNAs, which were downregulated after Sunitinib treatment while upregulated in metastasis tissues (Figure 1B). Notably, two potential candidate miRNAs (miRNA-452-5p and miRNA-605-5p) were selected on the basis of their involvement in RCC tumorigenesis by using OncomiR, an online resource for exploring miRNA dysregulation in cancer (Supplementary Figure A). As shown in Figure 1C and Supplementary Figure B, we used qRT-PCR to detect both miRNAs expression under Sunitinib treatment (5μM and 10μM) and finally selected out miRNA-452-5p as a validation target in OSRC-2 and SW839 cell lines.

**Figure 1.**
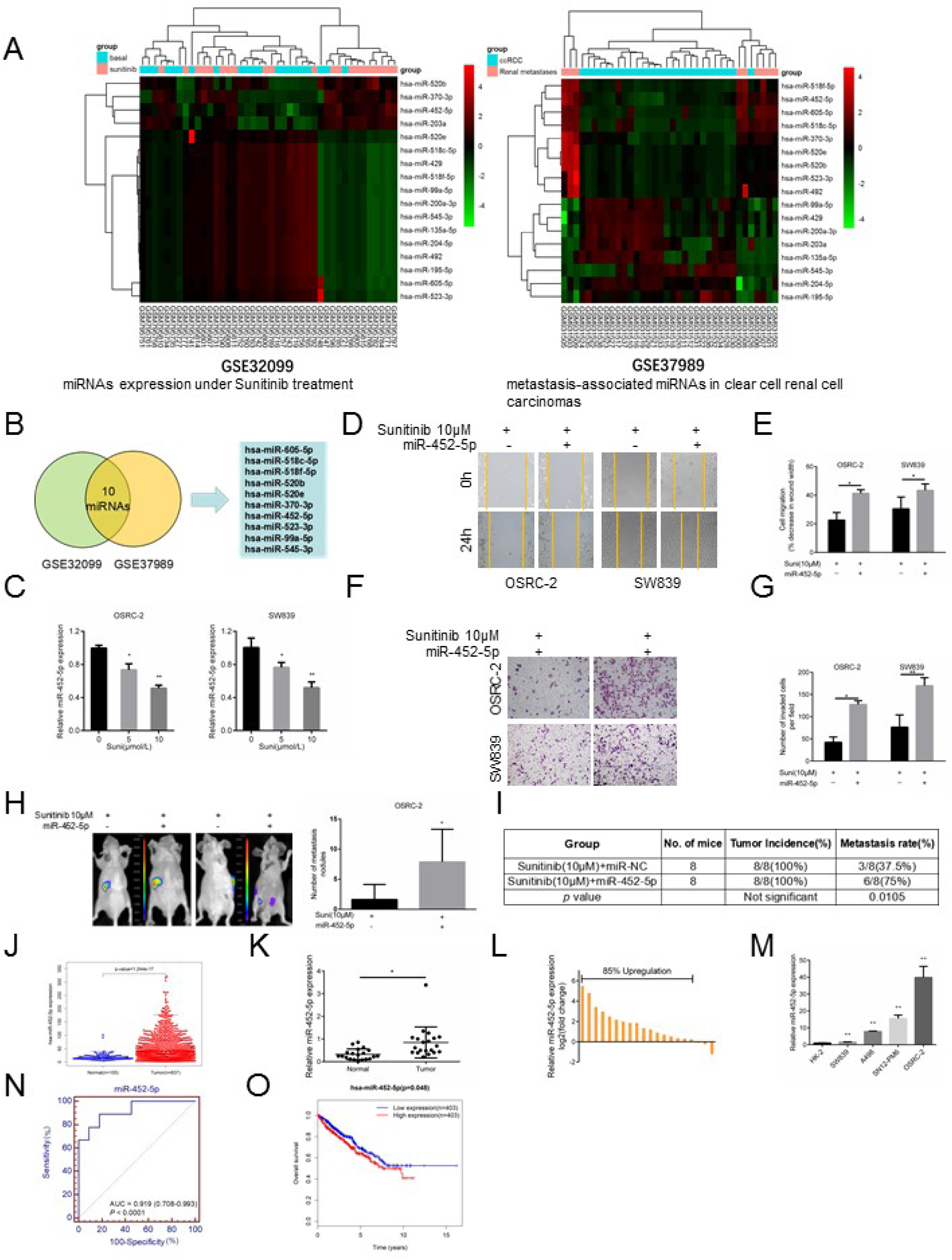
Sunitinib abrogates RCC cell invasion and metastasis via depressing miR-452-5p. A. Presented are heatmap of the most differentially expressed miRNAs in peripheral blood under Sunitinib treatment (GSE32099) and between tumor tissue and pair-matched normal tissues(GSE37989). B. The above TCGA analysis showed 10 miRNAs were significantly differentially expressed both in GSE32099 and GSE37989. C. qRT-PCR assays for miR-452-5p expression with 0, 5 and 10 μM Sunitinib treatment in OSRC-2 and SW839 cells. D-E. Representative micrographs of wound-healing assay and decrease in wound width was induced by transfection of miR-452-5p in OSRC-2 and SW839 cells versus miR-NC cells after treatment with 10 μM Sunitinib. F-G. Representative images and number of invasive cells per high-power field was induced by transfection of miR-452-5p in OSRC-2 and SW839 cells versus miR-NC cells after treatment with 10 μM Sunitinib. H. Orthotopic xenograft animal models were generated using miR-452-5p or miR-NC in OSRC-2 cells and mice treated with 10 μM Sunitinib. Presented are representative images (left) of abdominal metastasis viewed by IVIS system in each group 4 weeks after the orthotopic xenograft transplantation (n=8) and Quantitation of metastasis nodules shown at right. I. Incidence of metastasis in orthotopic xenografts after 4 weeks. J. TCGA cohort analysis of the differentially expressed levels of miR-452-5p in RCC tumor samples and pair-matched normal tissues. Each dot represents one sample. K. Comparison of miR-452-5p expression in 20 paired RCC tumor tissues and adjacent normal tissues via qRT-PCR. L. Relative miR-452-5p expression levels in RCC samples are presented as fold change=2^(ΔCt normal-ΔCt tumor)^ of tumor versus matched normal tissues, 85% of which was upregulated in tumor tissues than adjacent normal tissues. M. miR-452-5p expression in a series of RCC cell lines (SW839, A498, SN12-PM6 and OSRC-2) and human normal renal tubular epithelial cell line HK-2. N. ROC analysis to assess the specificity and sensitivity of miR-452-5p to differentiate between RCC and normal tissues. O. Kaplan–Meier analyses of the correlations between miR-452-5p expression level and overall survival of 806 patients with RCC through TCGA analysis. The median expression level was used as the cutoff. Data shown are mean±S.D. *P<0.05, \*\**P*<0.01

To confirm whether Sunitinib modulated miR-452-5p to impact its therapeutic effect, we first established RCC cell lines with miR-452-5p mimic, including OSRC-2 and SW839 (Supplementary Figure C). Transwell invasion assay suggested that miR-452-5p mimic-transduced both cell lines contributed to more invasiveness compared with cells with miR-NC group (Supplementary Figure D and E). Consistently, wound healing assay also indicated that miR-452-5p overexpression saliently enhanced cell migration than miR-NC group (Supplementary Figure F and G). Next, we prepared both cells lines with miR-452-5p mimic and miR-NC and then treated them with 10μM Sunitinib for 24h. As shown in Figures 1D and E, cells with miR-452-5p mimic performed more migratory capability compared with cells with control using wound-healing assay. Similarly, cells transfected with miR-452-5p mimic owned more invasive ability than those with control group (Figure 1F and G). Both pro-metastatic phenotype above elicited that miR-452-5p was potential one of the key mediators to influence Sunitinib efficacy.

To further confirm that miR-452-5p might act as a metastatic-promoting miRNA, we then established miR-452-5p with Sunitinib in a xenograft model. OSRC-2 cells with firefly luciferase expression were transfected with miR-NC or miR-452-5p under Sunitinib 10μM treatment. The stable clones were injected into left renal capsule of nude mice and metastatic sites were measured. As shown in the Figure 1H-I, Sunitinib blunted metastatic sites, which was partially reversed by miR-452-5p. Taken together, the results above suggested that miR-452-5p might be involved in Sunitinib repressing RCC invasion and metastasis.

### miR-452-5p is associated with poor prognosis of RCC

To further understand the expression and clinical relevance of miR-452-5p in RCC, the tumor development and survival outcome data were collected from The Cancer Genome Atlas (TCGA) database (http://cancergenome.nih.gov/). The results illustrated that the expression of miR-452-5p was pronouncedly increased in RCC tumor tissue compared to normal tissues (p<0.001) (Figure 1J and Table S4). In addition, qRT-PCR was performed in renal tumors and paired non-cancerous tissues from 20 RCC patients. The results demonstrated that the expression of miR-452-5p was substantially increased in RCC compared to adjacent non-cancerous tissues (p<0.05) (Figure 1K and L). As expected, the expression of miR-452-5p was remarkably expressed in various RCC cell lines, involving SW839, A498, SN12-PM6 and OSRC-2 compared to normal adult human kidney HK-2 cells (p<0.01) (Figure 1M). Furthermore, receiver operating characteristics (ROC) analysis revealed that miR-452-5p might serve as a useful biomarker for discriminating RCC from normal tissues with an area under the ROC curve (AUC) of 0.919 (95% CI, 0.708–0.993; Figure 1N). Next, we analyzed the association between miR-452-5p expression and the clinicopathological characteristics from 102 RCC patients (Table S5). Correlation regression analysis showed that high expression of miR-452-5p was significantly correlated with metastasis (p=0.043). We further performed univariate and multivariable logistic regression models to analyze the correlation of miR-452-5p levels with overall survival of RCC patients. Patient characteristics were provided in Table S6. Univariate analysis proved that a higher miR-452-5p expression (hazard ratio, HR=1.56; 95% confidence interval, CI=1.08–2.25; p=0.018) and metastasis (HR=1.64; 95%CI=1.25–1.86; p=0.036) were clearly correlated with overall survival. Multivariate analysis indicated that a higher miR-452-5p expression (HR=1.58; 95%CI=1.07–2.31; p=0.020) were markedly associated with overall survival. In addition, Kaplan–Meier survival analysis indicated that RCC patients with the lower levels of miR-452-5p had longer overall survival than those with the higher levels of miR-452-5p (p<0.05) (Figure 1O and Table S7).

In conclusion, the above data indicated that higher level of miR-452-5p was obviously correlated with poor prognosis of RCC.

### miR-452-5p-inhibition silences the development of pro-metastatic phenotype *in vitro*

Previously we proved that miR-452-5p facilitated cell migration and invasion in OSRC-2 and SW839 cells. To identify and characterize the biological function of miR-452-5p in RCC development, anti-miR-452-5p or anti-miR-NC were transfected into both RCC cells to investigate the anti-migratory and anti-invasive role of silenced miR-452-5p in renal cancer (Figure 2A). Compared to control group, anti-miR-452-5p could effectively inhibit migration and invasion in both RCC cells (Figure 2B-E). Furthermore, locked nucleic acid technology (LNA) was used to both cell lines to interrupt the expression of miR-452-5p (Figure 2F). The invasion and migration assay suggested the same tendency, which coincided with previous outcomes (Figure 2G-J).

**Figure 2.**
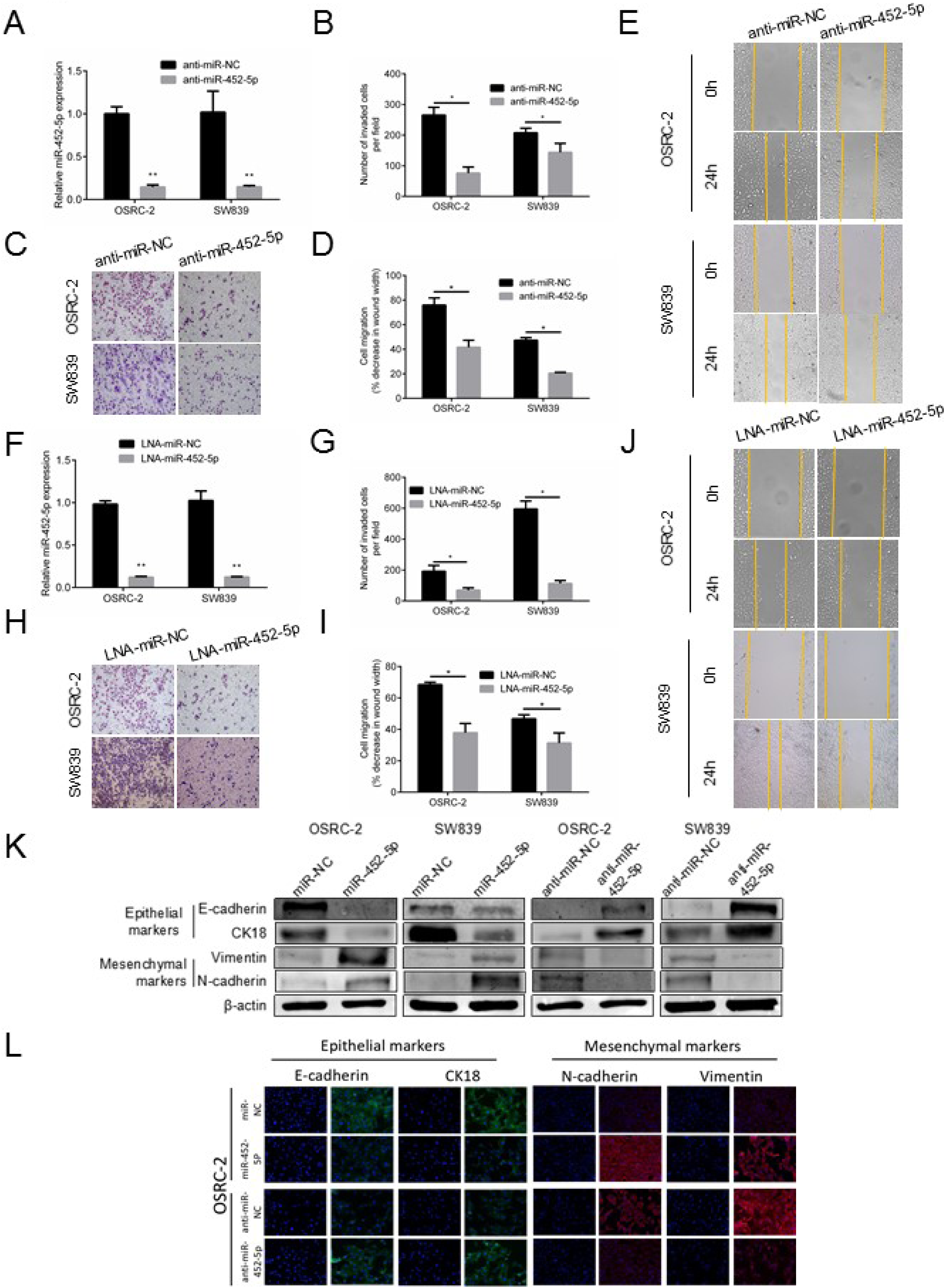
miR-452-5p promotes the development of pro-metastatic phenotype in vitro. A. qRT-PCR assays for anti-miR-452-5p and anti-miR-NC levels in OSRC-2 and SW839 cells, respectively. B-E. Wound-healing assay and transwell migration and invasion assay after the transfection with anti-miR-452-5p in OSRC-2 and SW839 cells when compared to that of anti-miR-NC cells. F. qRT-PCR assays for miR-452-5p and miR-NC levels in OSRC-2 and SW839 cells, respectively. G-J. Wound-healing assay and transwell migration and invasion assay after the transfection with miR-452-5p in OSRC-2 and SW839 cells when compared to that of miR-NC cells. K. Western blot analysis for EMT molecular markers (E-cadherin, CK-18, N-cadherin and Vimentin) protein levels of miR-452-5p comparing to miR-NC and anti-miR-452-5p comparing to anti-miR-NC transfection in OSRC-2 and SW839 cell lines. L. Immunofluorescence staining of EMT molecular markers (E-cadherin, CK-18, N-cadherin and Vimentin). Cells were respectively transfected with miR-NC, miR-452-5p, anti-miR-NC and anti-miR-452-5p. Data shown are mean±S.D. *P<0.05, \*\**P*<0.01.

It is widely recognized that EMT was implicated in cell invasion and metastasis. Here, we confirmed that miR-452-5p could induce EMT by immunofluorescence (IF) and western blot analysis of EMT marker expression. Using WB, we substantiated that miR-452-5p mimic-transduction decreased epithelial markers (E-cadherin and CK-18) expression while increased mesenchymal markers (N-cadherin and Vimentin) expression in OSRC-2 and SW839. Conversely, inhibition of miR-452-5p had the opposite effect, which epithelial markers (E-cadherin and CK-18) expression was increased yet mesenchymal markers (N-cadherin and Vimentin) expression was decreased in both cells (Figure 2K). Moreover, IF also showed that overexpression of miR-452-5p suppressed E-cadherin and CK-18 staining, while induced N-cadherin and vimentin expression in OSRC-2 cells. In contrast, E-cadherin and CK-18 expression was enhanced, whereas N-cadherin and vimentin expression was inhibited, where anti-miR-452-5p was transfected compared with the parental cells in OSRC-2 (Figure 2L).

Taken together, the above data identified that miR-452-5p functioned as a metastasis-promoting miRNA in RCC.

### miR-452-5p level is maximized by P65 through directly binding its promoter

To investigate the mechanism responsible for the up-regulation of miR-452-5p in RCC, we predicted 4 potential transcription factors, involving AR, P65 (RELA), HOXD9 and POU2F2, in the promoter region of miR-452-5p using PROMO, and using two miRNA target-predicting algorithms including Target Scan and miTar to narrow the candidates (Figure 3A). Then, UALCAN, an interactive web resource for analyzing cancer transcriptome data, was used to assess the above 4 transcription factors. Especially, AR and HOXD9 were ruled out due to their lower expression in RCC tissues than in normal renal tissues (Supplementary Figure H).

**Figure 3.**
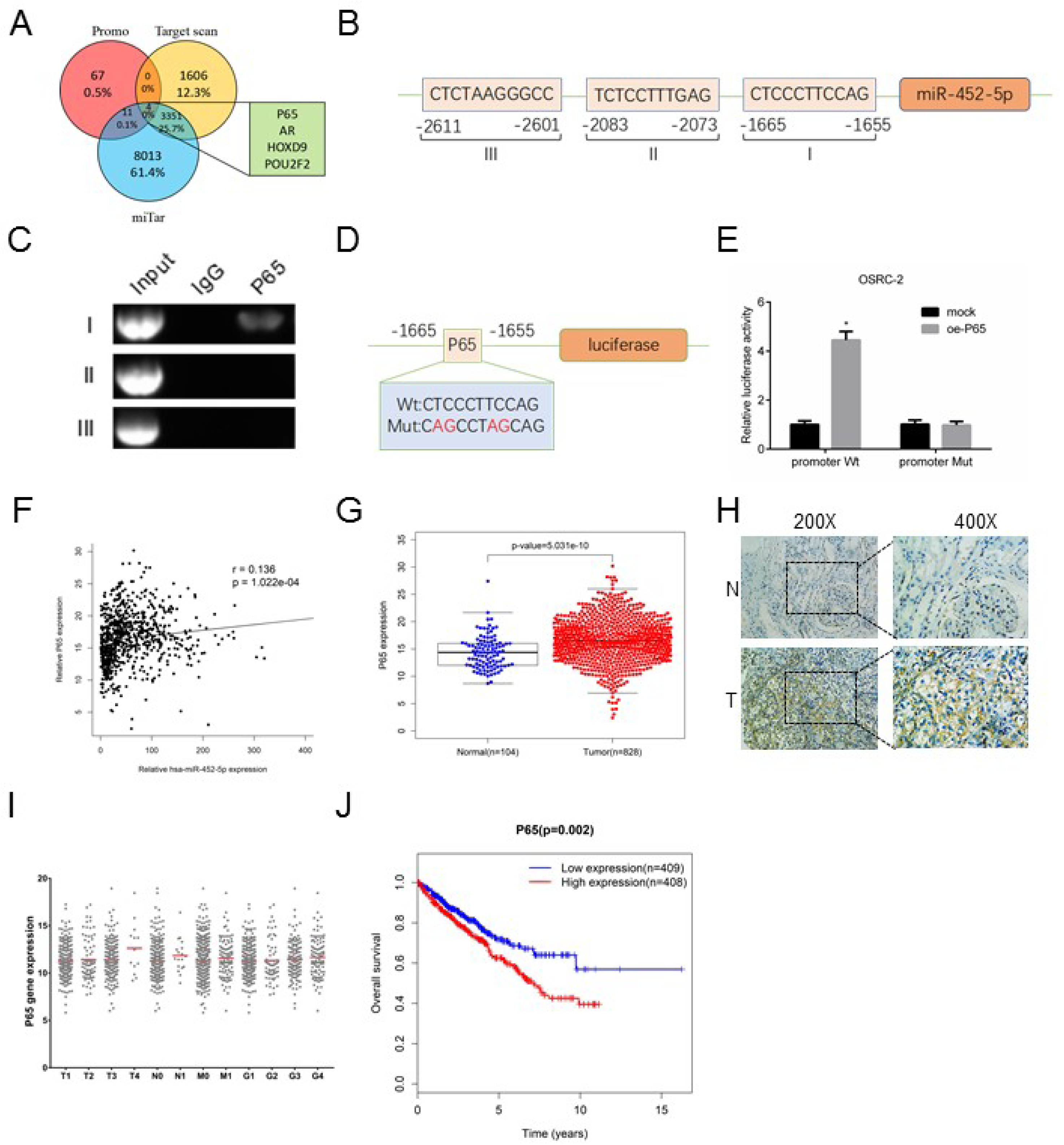
miR-452-5p level is maximized by P65 through directly binding its promoter. A. The schematic illustration of the proposed model depicting of 4 putative upstream transcription factors of miR-452-5p from 3 prediction databases, Promo, TargetScan and miTar. B. Bioinformatic analysis of potential P65 binding sites in miR-452-5p promoter. C. Lysates of OSRC-2 cells were subjected to ChIP assay and amplified by PCR reaction. D. The schematic illustration of P65-mut in the miR-452-5p promoter. E. Luciferase reporter assays in OSRC-2 cells. F. miR-452-5p expression and corresponding P65 expression in RCC and normal renal samples from RCC TCGA dataset. G. P65 expression in RCC and normal renal samples from RCC TCGA dataset. H. IHC staining revealed the level of P65 expression in RCC tissues (×200, × 400) versus adjacent normal tissues. I. P65 expression in different TNM stage of RCC samples from RCC TCGA dataset. J. P65 expression and patients’ survival time from RCC TCGA dataset. Data shown are mean±S.D. *P<0.05.

Next, we applied PROMO to determine the two transcription factors (P65 and POU2F2) binding sites of miR-452-5p within 3000 bases upstream from the transcriptional start site (Figure 3B and Supplementary Figure I). ChIP assay elucidated that P65 could bind to the promoter of miR-452-5p, while POU2F2 failed to bind (Figure 3C and Supplementary Figure J). Furthermore, luciferase assay confirmed that P65 increased miR-452-5p promoter activity, yet, this tendency abolished in miR-452-5p promoter-mutant OSRC-2 cells (Figure 3D and E).

Interestingly, data from TCGA database confirmed positive correlation between P65 and miR-452-5p expression (p<0.001) (Figure 3F and Table S8). In parallel, P65 expression was high expressed in RCC tumor tissues than in normal tissues (p<0.001) (Figure 3G and Table S9). In addition, Immunohistochemical (IHC) staining of P65 protein level indicated a higher expression in RCC tissues than in adjacent normal tissues (Figure 3H). Notably, TCGA database also showed higher P65 levels were obviously correlated with worse TNM stage (Figure 3I and Table S10). Kaplan–Meier survival analysis certified that patients with higher P65 levels had poor overall survival time than those with lower P65 levels from TCGA database (p<0.05) (Figure 3J and Table S11).

In brief, these results above corroborated that P65 were a carcinogenic gene and might induced miR-452-5p transcriptional level via directly binding its promoter in RCC.

### SMAD4 is the target gene of miR-452-5p and associated with good prognosis of RCC

To further dissect the mechanism underlying miR-452-5p modulating induction of RCC metastasis, we searched for potential downstream genes of miR-452-5p through four different miRNA target-predicting algorithms including TargetMiner, miRTarBase, miRWalk and miRTar, then focused on the one possible candidate target gene SMAD4 (Figure 4A). TCGA data sets suggested SMAD4 was low expressed in RCC tissues compared with normal tissues (p<0.05) (Figure 4B and Table S12). In particular, IHC staining of SMAD4 expression in RCC tissues and normal tissues also elucidated the same tendency with the above results (Figure 4C). Additionally, Kaplan–Meier survival analysis proved that patients with lower SMAD4 levels had poor overall survival time than those with higher SMAD4 levels from TCGA database (p<0.05) (Figure 4D and Table S13). Data from TCGA database revealed that there is a negative correlation between miR-452-5p and SMAD4, in keeping with the notion that miRNAs negatively regulate gene expression (Figure 4E and Table S14). In order to verify the speculation, by a computational prediction of miRNA databases, we identified three putative binding sites of miR-452-5p with high complementarity in SMAD4 promoter region (Figure 4F). To identify whether Wt-A, Wt-B or Wt-C was functional, luciferase reporter assays was performed and results demonstrated that all of these predicted binding sites in the promoter region of SMAD4 were functional. (Figure 4G). As a former study described, SMAD7 might be a feasible downstream gene of SMAD4 [28]. Consequently, WB analysis was used to detect the SMAD4 and SMAD7 protein level. When we introduced miR-452-5p mimic into OSRC-2 and SW839 cell lines, the decrease in SMAD4 and SMAD7 protein was confirmed (Figure 4H). Conversely, both inhibited miR-452-5p and LNA-miR-452-5p markedly enhanced their protein level (Figure 4I and J). Histological analysis of SMAD7 protein status in RCC tissues and normal tissues also elucidated that SMAD7 was potently downregulated in RCC tissues but was nearly undetectable in normal renal tissues (Figure 4K).

**Figure 4.**
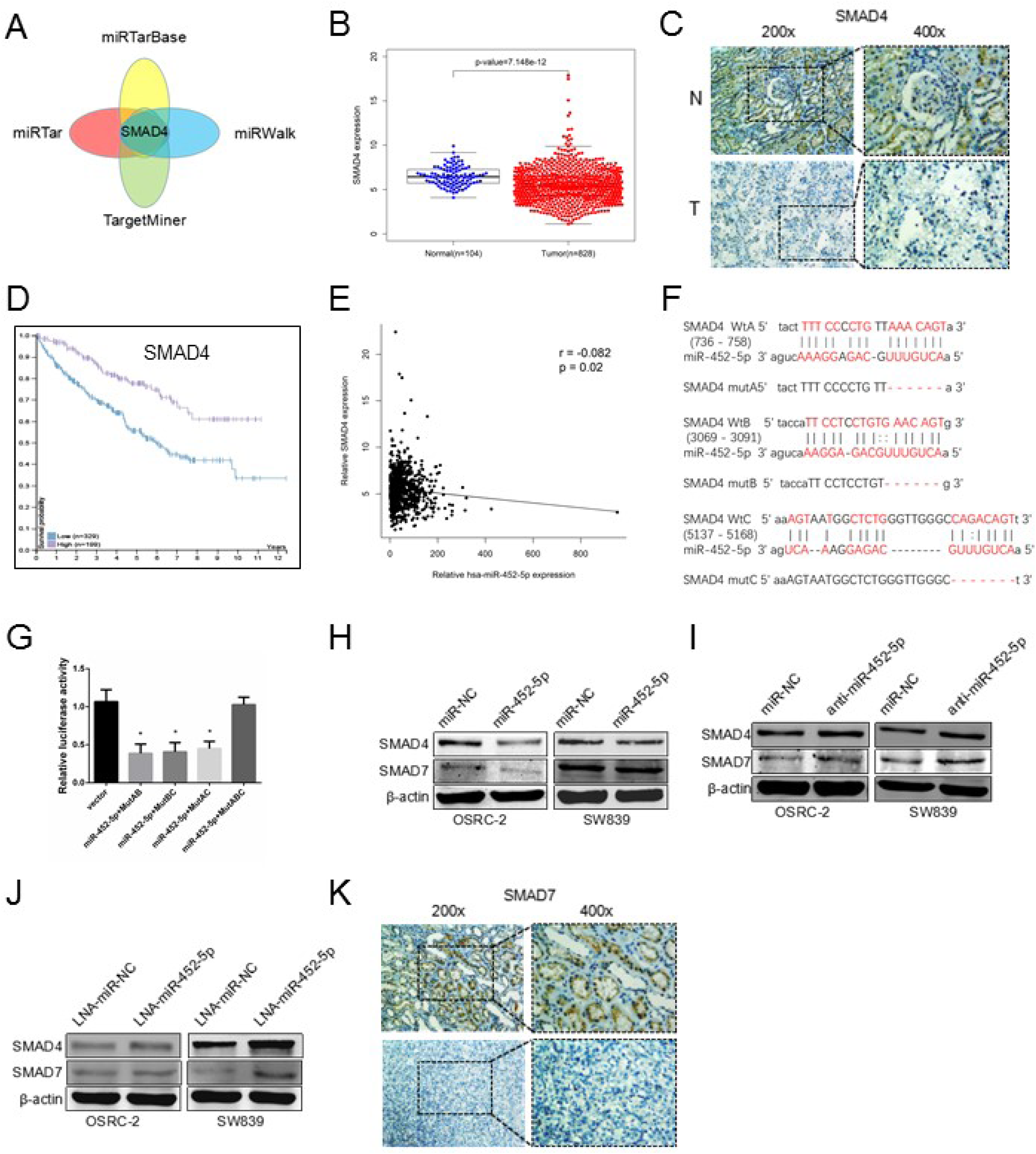
SMAD4 is the target gene of miR-452-5p and associated with good prognosis of RCC. A. The schematic illustration of the proposed model depicting of the putative downstream target gene, SMAD4, of miR-452-5p from 4 prediction databases, miRTarBase, miRTar, TargetMiner and miRwalk. B. SMAD4 expression in RCC and normal renal samples from RCC TCGA dataset. C. IHC staining revealed the level of SMAD4 expression in RCC tissues (×200, × 400) versus adjacent normal tissues. D. SMAD4 expression and paitients’ survival time from RCC TCGA dataset according to www.proteinatlas.org. E. miR-452-5p expression and corresponding SMAD4 expression in RCC and normal renal samples from RCC TCGA dataset. F. The schematic illustration of predicted binding site in the promoter in the 3’-UTRs of SMAD4 and sequences of the wild-type (SMAD4 WtA, WtB, WtC) and mutated 3’-UTR-Renilla luciferase reporters(SMAD4 mutA, mutB, mutC). G. Luciferase reporter assays in OSRC-2 cells. H-J. Western blot analysis for SMAD4 and SMAD7 protein levels of miR-452-5p comparing to miR-NC (H), anti-miR-452-5p comparing to anti-miR-NC (I) and LNA-miR-452-5p comparing to LNA-miR-NC (J) transfection in OSRC-2 and SW839 cell lines. K. IHC staining revealed the level of SMAD7 expression in RCC tissues (×200, × 400) versus adjacent normal tissues. Data shown are mean±S.D. *P<0.05.

Together, the data above revealed that miR-452-5p directly targeted SMAD4 and minimized SMAD4 and SMAD7 expression in RCC cells.

### SMAD4 recapitulates the effects of miR-452-5p in RCC cells

To examine whether SMAD4 could suppress cell invasion and metastasis in RCC, we performed gene set enrichment analysis with published gene array of metastatic RCC signatures (GSE12606), and results revealed that SMAD4 expression was negatively correlated with RCC cell migration, invasion and metastasis (Figure 5A). As previously described, we observed that miR-452-5p remarkably increased the cell migration and invasion of OSRC-2 and SW839 cells compared with mock. Importantly, an interruption approach with oe-SMAD4 partially reversed the effects of miR-452-5p on cell migration and invasion (Figure 5B and C). As we expected, WB confirmed that oe-SMAD4 recapitulated the effect of blocked SMAD4 and SMAD7 protein caused by miR-452-5p in both cell lines (Figure 5D).

**Figure 5.**
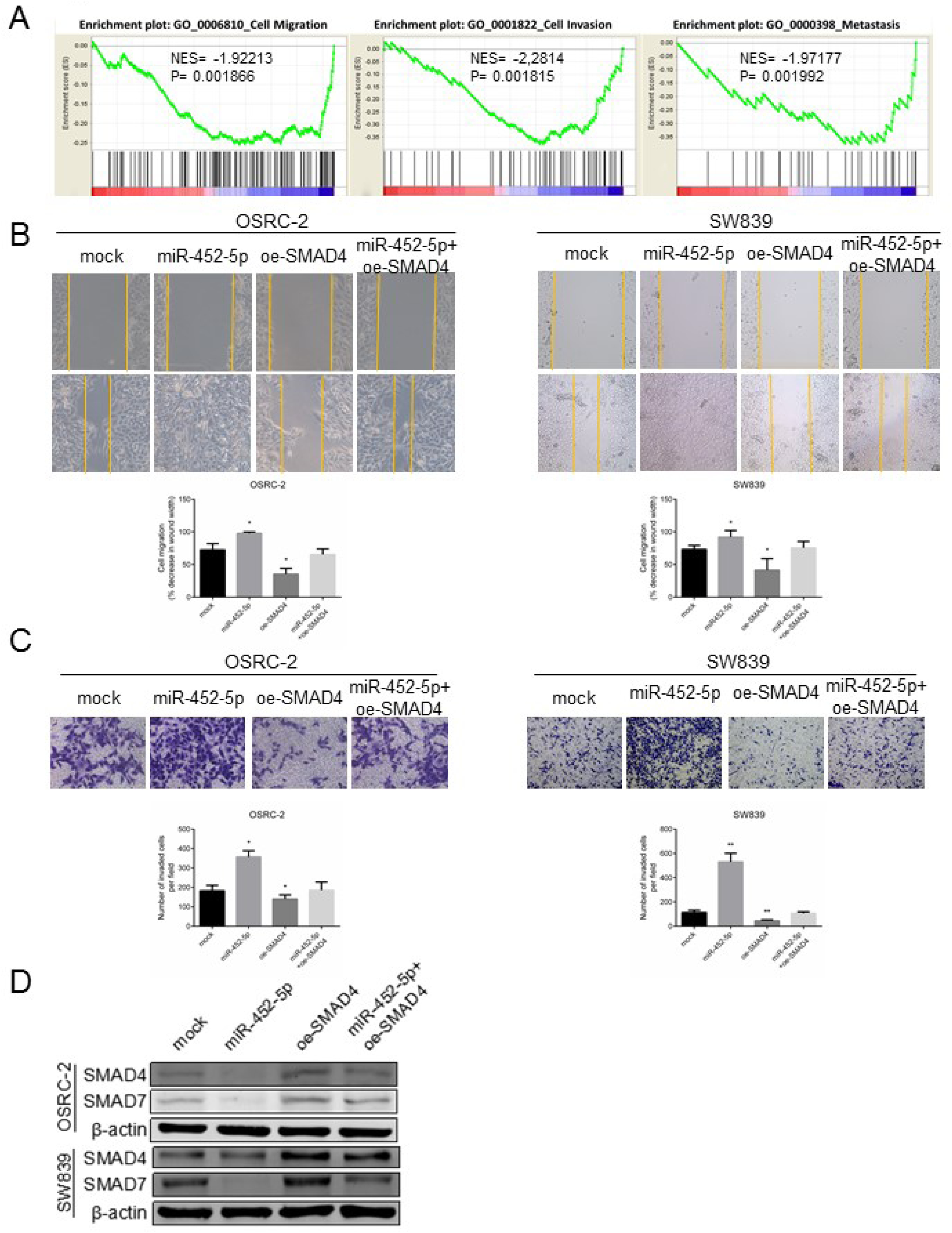
SMAD4 recapitulates the effects of miR-452-5p in RCC cells. A. GSEA of published gene array, GO:0006810, GO:0001822 and GO:0000398 databases referred to migration, invasion and metastasis related-gene signatures, respectively, of SMAD4 in RCC versus matched normal kidney tissue. NES, normalized enrichment score. B. Cell migration in OSRC-2 and SW839 cells were analyzed by wound-healing assay. Cells were respectively transfected with vector, miR-452-5p, oe-SMAD4 and miR-452-5p+oe-SMAD4. C. Cell invasion in OSRC-2 and SW839 cells were analyzed by transwell assays. Cells were respectively transfected with vector, miR-452-5p, oe-SMAD4 and miR-452-5p+oe-SMAD4. D. The protein levels of SMAD4 and SMAD7 in OSRC-2 and SW839 cells were assessed by Western blot analysis. Respectively, cells were transfected with vector, miR-452-5p, oe-SMAD4 and miR-452-5p+oe-SMAD4 for 48 hours before detection. Data shown are mean±S.D. *P<0.05, \*\**P*<0.01.

Together, these results above suggested that miR-452-5p promoted RCC cell invasion and metastasis through suppressing SMAD4.

### miR-452-5p promotes RCC metastasis through targeting SMAD4 *in vivo*

To further validate that miR-452-5p might act as a tumor inducer *in vivo*, we inoculated different clones of OSRC-2 cells. In this model system, sh-miR-452-5p cells as well as its control cells were inoculated into the left kidney capsule of xenograft. As shown in Figure 6A, a dramatic induction of luciferase expression in tumors of both groups was detected by in vivo imaging system (IVIS) as early as the 2th week. Figures 6B and C showed promotion of tumor metastases in the sh-miR-NC group compared with the sh-miR-452-5p group after 4 weeks (Figures 6B and C). Furthermore, sh-miR-452-5p attenuated lung, liver, spleen and right renal metastases (Figure 6D). In parallel, IHC staining substantiated SMAD4 and SMAD7 protein levels of sh-miR-452-5p group increased compared with sh-miR-NC group in renal tumor tissues from nude mice (Figure 6E). Conversely, transfection of miR-452-5p into OSRC-2 cells led to sufficiently enhanced metastasis of orthotopic xenograft tumors, whereas oe-SMAD4 into miR-452-5p upregulated cells mostly abolished this induction (Figure 6F-H). Together, results from Figure 6 illustrated that SMAD4-dependent miR-452-5p functioned as a critical tumor-metastasis promoter in RCC.

**Figure 6.**
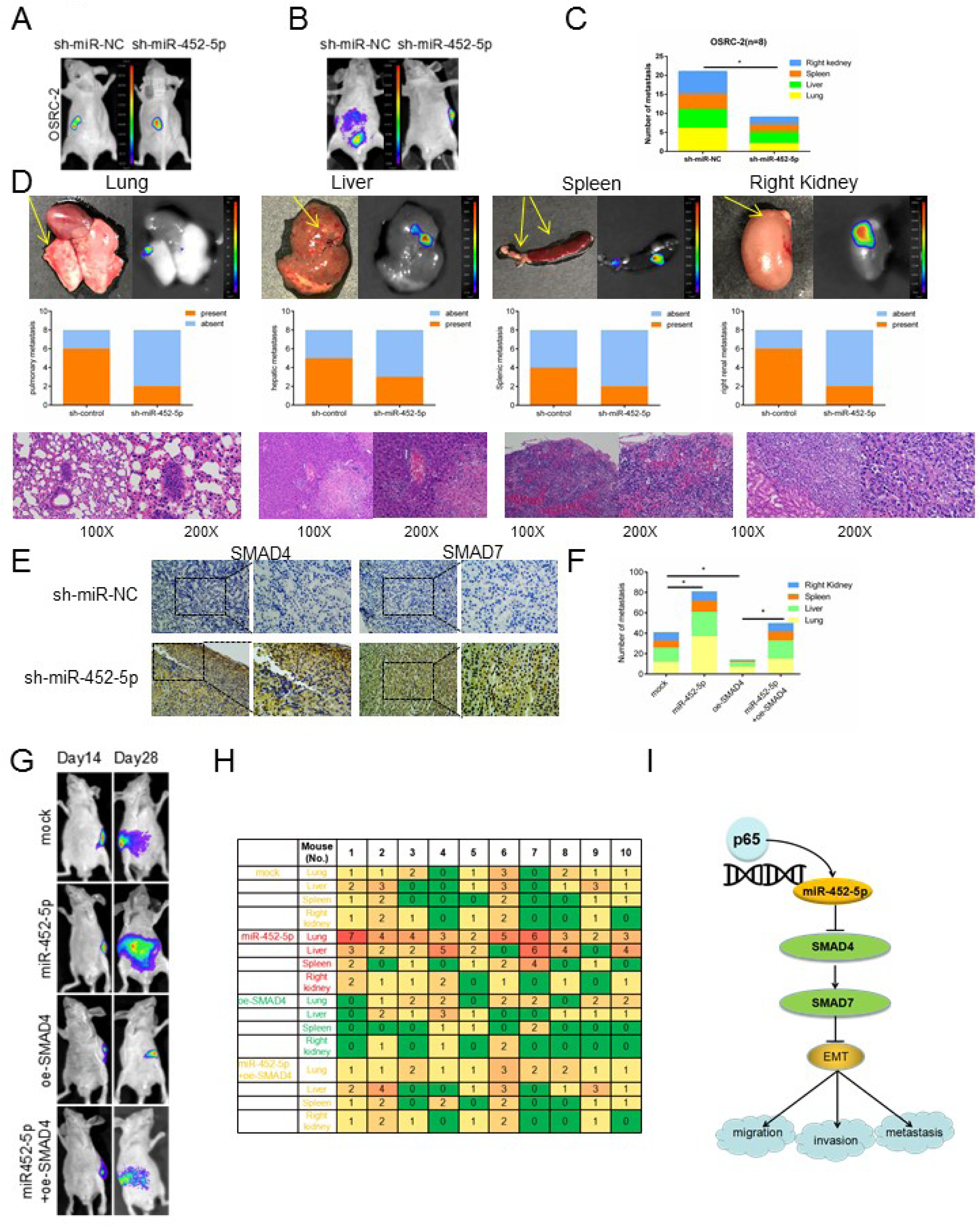
miR-452-5p promotes RCC metastasis through altering SMAD4 in vivo. A. Representative images of mice viewed by IVIS system in sh-miR-452-5p and sh-miR-NC group 2 weeks after left renal capsule injections (n=8). B. Representative images of abdominal metastasis viewed by IVIS system in each group 4 weeks after the orthotopic xenograft transplantation (n= 8). C. Presented are number of metastasis nodules in nude mice from the 7- to 8-week-old groups of sh-miR-452-5p and sh-miR-NC group. D. The animals were euthanized 4 weeks later for primary RCC and metastases detection by bioluminescent imaging, gross examination, and histological staining with haematoxylin and eosin (H & E). Quantitations were also shown. E. Representatives images of IHC staining of SMAD4 and SMAD7 inside or surrounding primary tumors, respectively. F-H. Orthotopic xenograft animal models were also generated using mock, miR-452-5p, oe-SMAD4 and miR-452-5p+ oe-SMAD4 transfected OSRC-2 cells. Presented are representative images (H) of abdominal metastasis viewed by bioluminescent imaging in each group 4 weeks after the orthotopic xenograft transplantation (n=10). Quantitations shown at right (F,H). I. A schematic for SMAD4/SMAD7/EMT Signals regulated by miR-452-5p. Data shown are mean±S.D. *P<0.05.

## Discussion

In our study, we first reported that miR-452-5p, induced by P65, was a potential therapeutic target for Sunitinib and associated with mRCC poor prognosis. MiR-452-5p in RCC samples and cell lines was high expressed compared with that in surrounding non-tumor tissues as well as HK2 normal cells. Moreover, P65 could directly bind to the promoter of miR-452-5p and transcriptionally induce miR-452-5p expression. Consistently, miR-452-5p elevated cell migration and invasion in RCC cell lines and promoted RCC progression through targeting SMAD4/SMAD7/EMT signals. All these results supported the conclusion that miR-452-5p acted as a tumor inducer and a metastasis-promoting miRNA in mRCC.

Previously, some reports demonstrated that miR-452-5p might function as a tumorigenesis-promoting in hepatocellular carcinoma and contribute to the docetaxel resistance of breast cancer cells [12; 49]. On the other hand, it has also been reported that expression of miR-452-5p inhibited metastasis in osteosarcoma, non-small cell lung cancer, and down-regulation of miR-452-5p is associated with adriamycin-resistance in breast cancer cells [13; 20; 47]. The possible reason for the opposite role of miR-452-5p in various human cancers or under different treatment is distinguished. Importantly, the relationship between mRCC and miR-452-5p as well as its response to Sunitinib remained unknown. In this work, our studies reported that Sunitinib attenuated miR-452-5p to impact its therapeutic effect, and miR-452-5p enhanced the development of pro-metastatic phenotype in mRCC.

According to early reports, P65, the central component of NF-κB pathway, has been suggested to act as a transcription factor in various kinds of diseases [7; 17; 35; 50]. Dong Yang *et al.* identified that P65 induced miR-17 transcription by binding its promoter elements to regulate proliferation of vascular smooth muscle cells under inflammation [45]. Rezapour S *et al.* reported that P65 was constitutively activated in colorectal cancer [35]. Other reports proved that P65 might serve as an activating transcription factor in several types of human cancers [19; 23]. Our conclusions supported these findings above that P65 could directly bind to the miR-452-5p promoter and thus transcriptionally induce miR-452-5p expression. Conversely, it was also reported that P65 could attenuate transcriptional activity under a certain condition. Raman P. NAGARAJAN *et al.* validated that P65 was able to inhibit the SMAD7 promoter activity [28], which, in some extent, was also coincided with our result that P65 induced miR-452-5p transcription, which targeted SMAD4 and repressed SMAD7 expression. Our study further illustrated that high P65 expression was obviously correlated with higher clinical TMN stage and contributed to poor prognosis in RCC patients. All these results concluded that P65 served as a tumor-inducer in RCC.

As a member of SMAD family, SMAD4 is a critical component of TGF-β signaling and gets involved in MAPK, CDK and PI3K signaling [29; 48]. As we all know, SMAD4 has been recognized as a tumor suppressor gene in a good deal of cancers, and recent study has presented that SMAD4 suppressed the progression of RCC by targeting various downstream genes [14; 24; 26; 44]. Moreover, another report suggested that the loss of SMAD4 repressed SMAD7 transcription leading to a loss of functional protein during renal inflammation and fibrosis [28]. Here, we elucidated that miR-452-5p directly targeted SMAD4, and as a downstream gene, SMAD7 was also repressed. Furthermore, SMAD7 has been reported to play a pivotal role in EMT, which finally contributed to tumor metastasis [36; 41; 43]. Our study presented a road map that miR-452-5p facilitated RCC invasion and metastasis through SMAD4/SMAD7/EMT Signals.

Although Sunitinib is the first-line treatment for mRCC, the clinical benefit of sunitinib is limited, and the vast majority of mRCC patients under sunitinib treatment ultimately develop disease progression because of the acquisition of resistance. Several studies reported that miRNAs play a crucial role in alterting the sensitivity to Sunitinib in multiple tumors, indicating that miRNAs are potential therapeutic targets for Sunitinib, and miRNA modulation combined with Sunitinib as a novel therapeutic strategy was under exploring [5; 10; 16; 34]. In our study, miR-452-5p was confirmed to determine the sensitivity to Sunitinib, and high expression of miR-452-5p was suggested to contribute to Sunitinib resistance. Therefore, it could be used to divide mRCC patients into different responsive groups to save both money and time for the non-responsive patients and enhance Sunitinib treatment efficacy through up-regulating the expression of miR-452-5p.

In this study, we conclude that Sunitinib efficacy is potentially connected with miR-452-5p, which acts as an efficient metastasis-promoter through SMAD4/SMAD7/EMT signals in RCC patients. This finding points to a novel therapeutic target to maximize Sunitinib efficiency in RCC, and helps us to better suppress RCC progression via targeting this newly identified signal pathway.

## Acknowledgements

This work was sponsored by the National Science Foundation of China (81602216 and 81671446), National Science Foundation of Shanghai (16ZR1426500), Shanghai Pujiang Program (16PJD037).

